# MINOTAUR: A platform for the analysis and visualization of multivariate results from genome scans with R Shiny

**DOI:** 10.1101/062158

**Authors:** Robert Verity, Caitlin Collins, Daren C. Card, Sara M. Schaal, Liuyang Wang, Katie E. Lotterhos

## Abstract

Genome scans are widely used to identify “outliers” in genomic data: loci with different patterns compared with the rest of the genome due to the action of selection or other non-adaptive forces of evolution. These genomic datasets are often high-dimensional, with complex correlation structures among variables, making it a challenge to identify outliers in a robust way. The Mahalanobis distance has been widely used for this purpose, but has the major limitation of assuming that data follow a simple parametric distribution. Here we develop three new metrics that can be used to identify outliers in multivariate space, while making no strong assumptions about the distribution of the data. These metrics are implemented in the R package MINOTAUR, which also includes an interactive web-based application for visualizing outliers in high-dimensional datasets. We illustrate how these metrics can be used to identify outliers from simulated genetic data, and discuss some of the limitations they may face in application.

## Introduction

Knowledge of the genetic architecture of biological traits—the number of loci that affect a phenotype, the magnitude of their effect, and their distribution across the genome—not only illuminates the evolutionary processes that shape genomes, but also has important implications for complex diseases (McCarthy and Hirschhorn 2008), conservation (Kohn et al. 2006; Allendorf et al. 2010; Funk et al. 2012), and breeding programs (Goddard et al. 2009; Varshney et al. 2009). With the advent of next-generation sequencing we now have the ability to examine genomes at a fine scale; and, as a result, we have identified a large number of genomic variants that are implicated in complex diseases (Carlson et al. 2004; Hindorff et al. 2009) and adaptation to the local environment (Savolainen et al. 2013). This wealth of data is likely to yield new insights, but it also brings with it the challenge of extracting the relevant signal from noisy, complex, multi-dimensional data sets. This is perhaps one reason why most of the variants detected so far have only managed to explain a very small proportion of the observable phenotypic variation (Yang et al. 2010; Brachi et al. 2011).

The preferred method for detecting genomic variants is via genome scans. There are many different approaches toward scanning genomes, but all are based on the same premise: that the loci of interest to the investigator are likely to be statistical outliers when compared with the rest of the genome. The particular choice of statistic will depend on the question being asked and the experimental design, and may include one or more statistics from the following categories: tests for genetic differentiation (Lotterhos & Whitlock 2014; Hoban et al. *in revision*), scans for strong positive selection and/or selective sweeps (Hohenlohe 2010; Vatsiou et al. 2016), genome-wide association studies for phenotype-associated loci (GWAS, reviewed in Carlson et al. 2004 and McCarthy et al. 2008), linkage mapping for quantitative trait loci (QTL, Savolainen et al. 2013), genetic-environment associations (reviewed in Rellstab et al. 2015), and scans for differentially expressed genes (Wang et al. 2009). A number of different genome-scan test statistics may be calculated for a single genomic dataset and these are usually examined one-at-a-time (i.e., in univariate analyses). Some test statistics may be highly correlated, while the power of other test statistics may vary for different regions of the genome depending on the details of selection, recombination, mutation, and migration rates (Tiffin and Ross-Ibarra 2014). Additionally, the power of different approaches may vary among species because of demographic history, and within a species because of sampling design (De Mita et al. 2013; de Villemereuil et al. 2014; Lotterhos and Whitlock 2015). Finally, loci with intermediate probabilities of detection will often exhibit the highest variance in results from genome scans (Lotterhos et al. *in review*).

Given the complex evolutionary histories of most species, it is doubtful whether any single statistic can fully capture the genomic signal of interest in the majority of cases (Verity and Nichols 2014). Furthermore, the uncertainty in demographic history, coupled with the variation in statistical outcomes in different scenarios, makes it difficult to know which statistics have the greatest power to detect selection and which have the highest false positive rates. These issues point to a need for composite, multivariate outlier methods that integrate information across multiple test statistics.

Multivariate methods have been utilized extensively in many biological applications, although in application to genome scans the power of the multivariate approach for detecting outliers has not yet been fully evaluated. Because some dimension reduction methods such as Principal Component Analysis rely on assumptions about the data that may be unjustifiable in the context of genome scans (O’reilly et al. 2012), these methods are not ideally designed for the identification of multivariate outliers (Pattterson et al. 2006). Some GWAS analyses have successfully employed multivariate approaches to identify genetic associations with multiple phenotypes (O’reilly et al. 2012; Galesloot et al. 2014). Additionally, multivariate approaches have also been used in GWAS metaanalysis to simultaneously consider multiple genetic or phenotypic variables (reviewed in Evangelou and Ioannidis 2013). It is evident, however, that more opportunities exist for the use of multivariate approaches in outlier detection than are currently being capitalized on.

While there are dedicated software tools for calculating a variety of test statistics, there does not currently exist a unified platform for the filtering, visualization, and integration of test statistics in multivariate space. Here we describe a new R package called MINOTAUR (<u>M</u>ultivariate v<u>I</u>sualisatio<u>N</u> and <u>O</u>u<u>T</u>lier <u>A</u>nalysis <u>U</u>sing <u>R</u>) built specifically for this purpose. This software package - initiated during a hackathon for population genetics in R (https://github.com/NESCent/r-popgen-hackathon) - provides functions for detecting outliers in multivariate space alongside procedures to manipulate, summarize, and visualize these data. The R software environment (R Core Team 2015) is free, open-source, and hosts a large collection of tools for statistical analysis, making it the ideal host for the development and uptake of such a platform. Furthermore, because data visualization is an important part of verifying and identifying outliers, the R Shiny and Shiny Dashboard environments (Chang 2015; Chang et al. 2016) have been employed to provide MINOTAUR users with an interactive interface that streamlines the process of data input, statistical analysis, and graphical exploration. Together, these tools have the potential to increase the efficiency with which the results of genome scans are interrogated.

## Approaches to identifying multivariate outliers

In the MINOTAUR package we implement four composite measures that can be used to integrate information over multiple univariate statistics: the Mahalanobis distance, harmonic mean distance, nearest neighbor distance, and kernel density deviance. We developed the latter three measures, which are related to Mahalanobis distance but make no strong assumptions about the parametric form of the data, meaning they can be applied to multivariate statistics that have complex correlated or even multimodal distributions. Some of these measures are heavily influenced by the distance of points from the multivariate centroid (Mahalanobis and harmonic mean distance) while others are mainly influenced by the sparseness of points in the local vicinity (nearest neighbor distance and kernel density deviance), and so we would expect the measures to behave differently from one another, and to vary in their behavior depending on the data at hand.

The calculation of these composite measures has been optimized for genome-scale data by using precompiled routines, written in C++ and integrated into R using the package Rcpp (Eddelbuettel and Francois 2011; Eddelbuettel 2013). Several packages devoted to multivariate statistics that may be appropriate for genome-scale data already exist in R (see Supplementary Table 1), and thus users are free to utilize both existing statistical methods and the more targeted functions included within the MINOTAUR package.

*Mahalanobis distance.* The Mahalanobis distance is a multidimensional measure of the number of standard deviations that a point lies from the mean of a distribution. The Mahalanobis distance of a *d*-dimensional observation *x_i_* = (*x*_*i*1_, *x*_*i*2_,…, *x_id_*)*^T^* from a distribution of *N* variables with mean 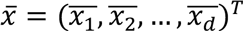 and covariance matrix *S* is defined as follows (Mahalanobis 1936):

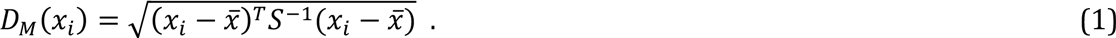

This distance differs from the ordinary Euclidean distance due to the correction for covariance among observations, making it a better distance measure for genome scan summary statistics because it does not assume that statistics are independent (i.e., Euclidean distance equals Mahalanobis distance when *S* is a diagonal matrix). However, this distance does make the assumption that points disperse smoothly from a single multivariate centroid, and so it will tend to perform poorly when observations have a complex or multimodal distribution.

*Harmonic mean distance.* In this context the “harmonic mean distance” of an observation *x_i_* refers to the harmonic mean of the distances between this point and all other points. The distance measure used here is the Euclidian distance normalized by multiplying by the inverse covariance matrix. This ensures that results are not dominated by a few statistics with a large spread, and also accounts for any potential correlation between statistics, analogously to the Mahalonobis distance. Mathematically we can define the harmonic mean distance as follows:

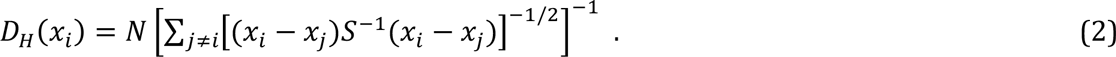

The harmonic mean is heavily influenced by small values, which in this context means local effects are amplified. However, more distant points also have some effect on the final value (unlike the nearest neighbor distance described below), and so the harmonic mean strikes a balance between local and global effects. This has some advantages in outlier detection, as observations that are both distant from the main mass of the data and have few neighbors in the local vicinity will tend to be outliers.

*Nearest neighbor distance.* The nearest neighbor distance of the observation *x_i_* gives the minimum distance between this point and any other point. As with the harmonic mean distance, we use the Euclidian distance normalized by the inverse covariance matrix. Mathematically we can write

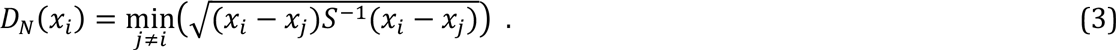

This statistic exclusively measures local effects, being largest when an observation is a long way from any other point. Because this distance is only based on two points (the focal point and its nearest neighbor), it is not influenced by the global distribution of the data, unlike the harmonic mean distance.

*Kernel density deviance.* Kernel density-based methods attempt to capture mathematically the distribution of the data as the sum of a number of simple parametric distributions. Here we apply these methods to identifying multivariate outliers, defined as those points with a low density of data around them in multivariate space. We assume a multivariate normal kernel *G*(*x_i_* | *x_j_*, *λ*^2^*S*) centered at the point *x_j_*, where λ is the bandwidth of the kernel, which is scaled in each dimension by the covariance matrix of the data. We then calculate the leave-one-out log-likelihood (Leiva-Murillo and Artés-Rodriguez, 2012) of the point *x_i_* as follows:

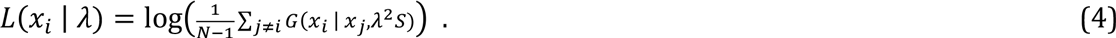

In other words, this is equal to the log-probability density of the point *x_i_* under the kernel density distribution constructed from all points apart from *x_i_*. Our final density-based measure is defined as follows:

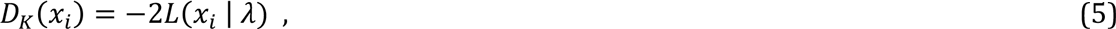

which is sometimes referred to as the Bayesian deviance. This will be large whenever the density of the point *x_i_* is low, and so the kernel density deviance can be thought of as a measure of the sparseness of points around the focal point.

One challenge when using kernel density methods is choosing an appropriate value for the bandwidth. Here we simply use the bandwidth for which the total deviance of all points is minimized, i.e.

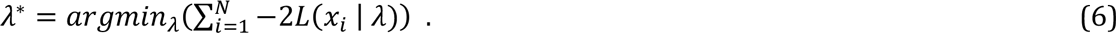

It can be shown that this is equivalent to the maximum-likelihood value of *λ* under the leave-one-out criterion. The value *λ*\* can be found using the MINOTAUR function `kernelDeviance`(), which takes a vector of bandwidths as input and returns the total deviance of each. This function can be used to search for the minimum value of *λ* manually, or via an optimization routine such as `optim`(). Users are also free to use any other bandwidth, entered manually, or in the absence of a user-defined bandwidth asimple method based on Silverman’s rule is implemented as a default (this assumes that data is normally distributed, and is a simple function of the standard deviation of the samples (Silverman 1986)).

## The MINOTAUR R package - an R Shiny graphical user interface for multivariate outlier analysis and visualization

The MINOTAUR package performs two main functions: (1) it calculates the compound multivariate outlier statistics described above and (2) it enables users to harness the interactive graphical power of the R Shiny environment to manipulate and visualize their data within the MINOTAUR graphical user interface (GUI). The GUI allows users to perform the former task with the click of a button; however, outlier identification can also be performed on the R command line using stand-alone functions available in MINOTAUR, if preferred. Directions for downloading and installing the package can be found at the end of this manuscript.

The MINOTAUR GUI is designed to streamline the process of genomic data analysis and outlier identification, taking users from data input to graphical output within a single platform. Distinct panels are used for each stage of the analysis, including data input and filtering, outlier detection via the methods described above, and plotting results (e.g., histograms, scatterplots, and Manhattan plots). An overview of the MINOTAUR GUI workflow is show in Figure 1.

**Figure 1.**
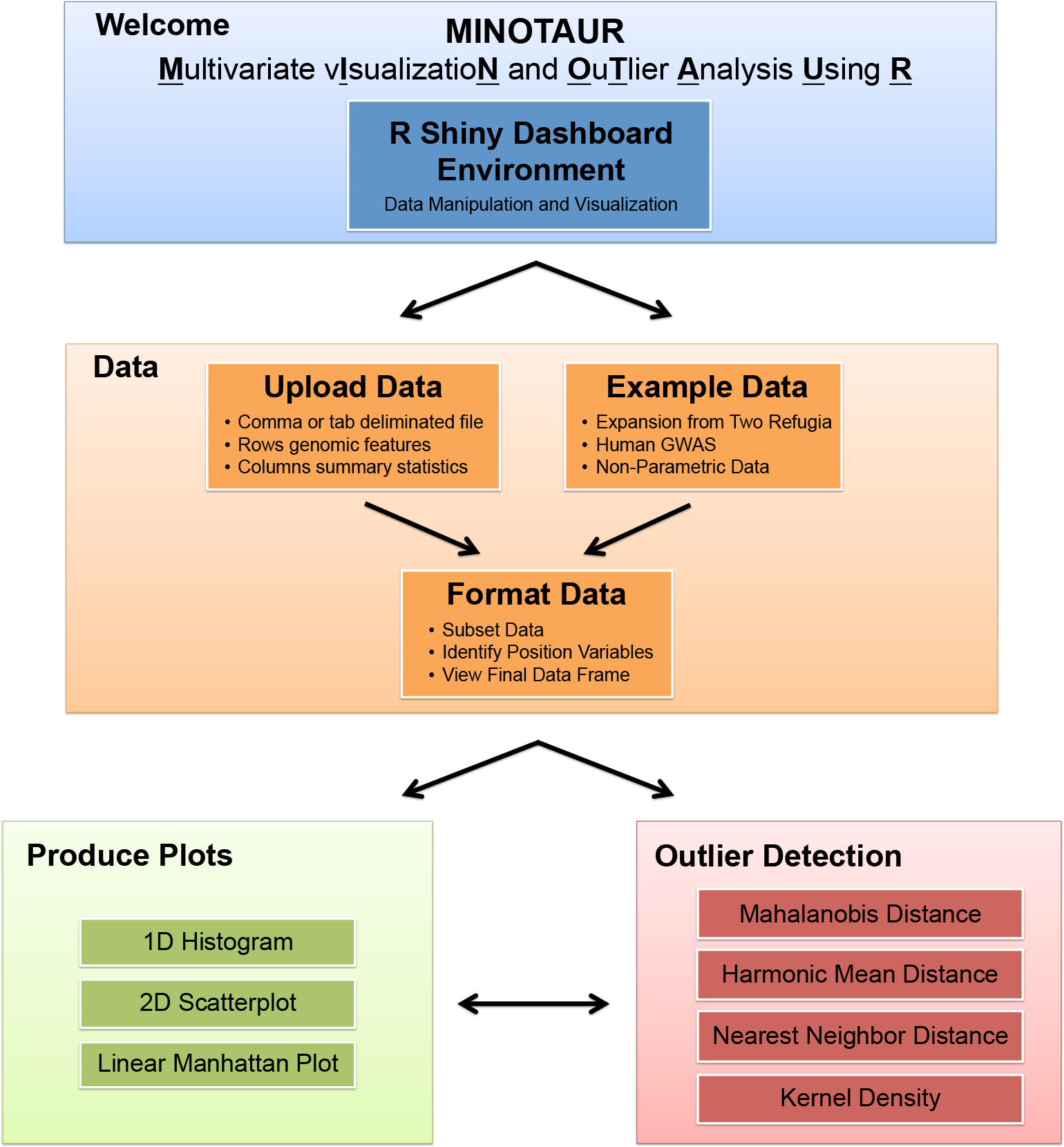
Graphical overview of the MINOTAUR GUI workflow.

In the *Data* panel, the MINOTAUR GUI allows users to either upload their own datasets or select among a set of four in-built example datasets. Data can be uploaded in a number of file formats, including comma-or tab-separated text files, and Rdata. Regardless of the file format, MINOTAUR expects all incoming datasets to be arranged in data frames, with each row representing a different genetic locus and each column representing a different univariate genome scan statistic (e.g., *F*_*ST*_, Tajima’s *D*, etc.) or other piece of locus-specific metadata (e.g., SNP identifiers, chromosomes/scaffolds and positions, etc.). Raw data objects can be filtered within the GUI, meaning, for example, that columns not related to outlier analysis can be dropped at an early stage.

Four example datasets are made available to users within the MINOTAUR package and GUI. The “HumanGWAS” dataset contains example output from an unpublished human Genome-Wide Association Study. The simulated “NonParametricInverse” and “NonParametricMultimodal” datasets each contain an example of nonparametric data, one with an inverse relationship (Figure 3) and one that is highly multimodal (Supplemental Figure S1). The “TwoRefSim” dataset contains population genetic data simulated under a model of expansion from two refugia (Lotterhos and Whitlock 2015). Note that the example datasets can also be accessed outside the GUI by running the `data`() command with the appropriate dataset name. For example, to load the “HumanGWAS” dataset, type `data`(`HumanGWAS`) and hit `ENTER`. To learn more about a dataset while in the R terminal, add a question mark before the dataset name to load the relevant Help page; for example, type `?HumanGWAS` and hit `ENTER`.

In the *Outlier Detection* panel, multiple univariate statistics can be integrated to produce the compound distance measures described above. These measures can be appended to the data frame and visualized interactively in the *Produce Plots* panel, which includes several submenus with useful plots for visualizing high-dimensional datasets, including Manhattan plots, 1D histograms and density-based 2D Scatterplots. The plotting methods are designed with large genomic datasets in mind; for example the `plot2d`() function included with the package calculates the density of points for a given bin size and shades bins according to the density of points within them, and then optionally adds user-supplied points (ideally a small subset of points, for example the outliers only) to the plot. Additional options allow users to log-scale statistics and control various other visual settings commonly used when plotting data in R (Figure 2).

**Figure 2.**
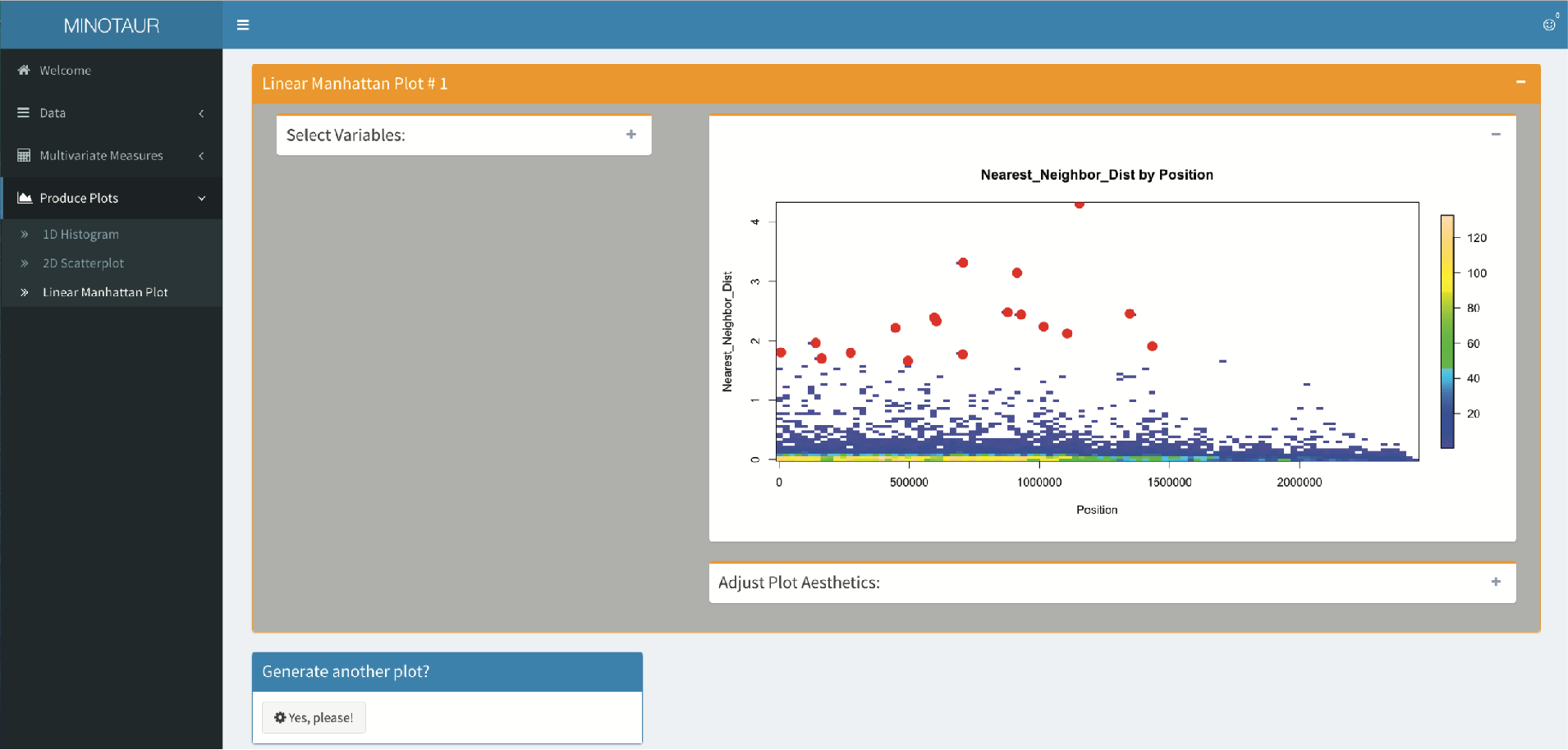
Screenshot of MINOTAUR GUI highlighting the overall interface and the ability to visualize multivariate distributions. The plot is a Manhattan plot of the nearest neighbor distance across loci for all traits in the “HumanGWAS” example dataset provided as part of MINOTAUR. The base scatter plot demonstrates the binned visualization, where the density of data in an area is apparent from the color. 99.5% percentile outliers are indicated with solid orange circles. Visualization menus have been collapsed to simplify the image. Additional plots can also be stacked below to enable comparisons across multiple plots (not shown).

## Example applications of multivariate outliers

*Evaluation of computational speed.* First, we evaluated the speed of calculating the four compound distance measures for datasets with increasing numbers of loci (rows) and univariate statistics (columns). For this example, variables were randomly generated from a multivariate normal distribution. Table 1 gives the “order” of complexity of these algorithms, together with measured run-times for a dataset composed of 50,000 loci and 10 variables (see Supplementary Table S2 for extended run-time analyses). Overall, the Mahalanobis distance is calculated in a matter of seconds, even with particularly large datasets. The harmonic mean distance, nearest neighbor distance, and kernel density deviance each scale approximately equally with increasing dataset sizes, though the maximum likelihood estimate of the ideal bandwidth for the latter measure can add significant computation time.

**Table 1.** Multivariate outlier detection methods implemented in MINOTAUR and associated computational run times. Computational complexity is given in “big O” notation, with *N* referring to the number of observations and *k* the number of statistics (dimensions). Run times were determined using an Apple iMac with a 3.1 GHz Intel Core i5 processor and 32 GB of RAM running Apple OSX 10.9.5 and R version 3.2.3. Note that for computation time the kernel density deviance includes both the maximum likelihood estimation of the optimal bandwidth and the density calculations based on the optimal bandwidth.

| Compound measure | Description | R Function | Computational complexity (big O notation) | Computation Elapsed Time for 50,000 loci & 10 variables (hh:mm:ss.ms) |
| --- | --- | --- | --- | --- |
| Mahalanobis distance | Distance from multivariate centroid | Mahalanobis() | $O(Nk^2)$ | 00:00:00.095 |
| Harmonic mean distance | Inverse-weighted distance from all other points | harmonicDist() | $O(Nk^2)$ | 00:04:13.620 |
| Kernel density deviance | Local density of points | kernelDist() | $O(Nk^2)$ | 01:40:03.600 |
| Nearest neighbor distance | Distance to nearest neighbor | neighborDist() | $O(Nk^2)$ | 00:04:07.020 |

*Example on simulated nonparametric distributions.* Some kinds of genomic data - for example gene expression data - may generate complex nonparametric distributions. Genes that have high expression in one environment may have low expression in another environment, while investigators may be interested in identifying genes that have moderate expression in both environments. To test the performance of the multivariate outlier statistics in nonparametric situations, we simulated two examples of nonparametric distributions.

In the first example, we simulated a distribution of two variables that follow an inverse relationship, with some additional noise. We used contour plots to visualize the different ways in which each of the compound distance measures changes over the twodimensional plane (Figure 3). In these plots, the darker red lines indicate less-significant values of the test statistic and lighter yellow lines indicate more-significant values of the test statistic. We also looked at two manually chosen points on the plane - indicated by a blue square and triangle - chosen to represent different sorts of outliers. The blue triangle would not be considered an outlier from the perspective of either onedimensional distribution despite being a clear outlier from the two-dimensional distribution, while the blue square would be considered an outlier in the first dimension but not the second. In this example, the nonparametric distribution affects the relative ability of the four statistics to identify each of these outliers (Figure 4). The blue triangle would not have the largest value (i.e., not be the most outlying point) by the Mahalanobis or the harmonic mean distance, while it would have the largest value by nearest neighbor distance or kernel density deviance. In contrast, the blue square has the largest value of the test statistic by all four methods.

In the second example, we simulated a highly multimodal distribution from a normal mixture model. In this example, it can be seen how the parametric assumption of the Mahalanobis distance fails to capture the complexity of the data (Supplementary Figure S1). In contrast to the previous example, the harmonic mean distance behaves similarly to the kernel density deviance, and nearest neighbor distance has the most complex contour landscape.

**Figure 3.**
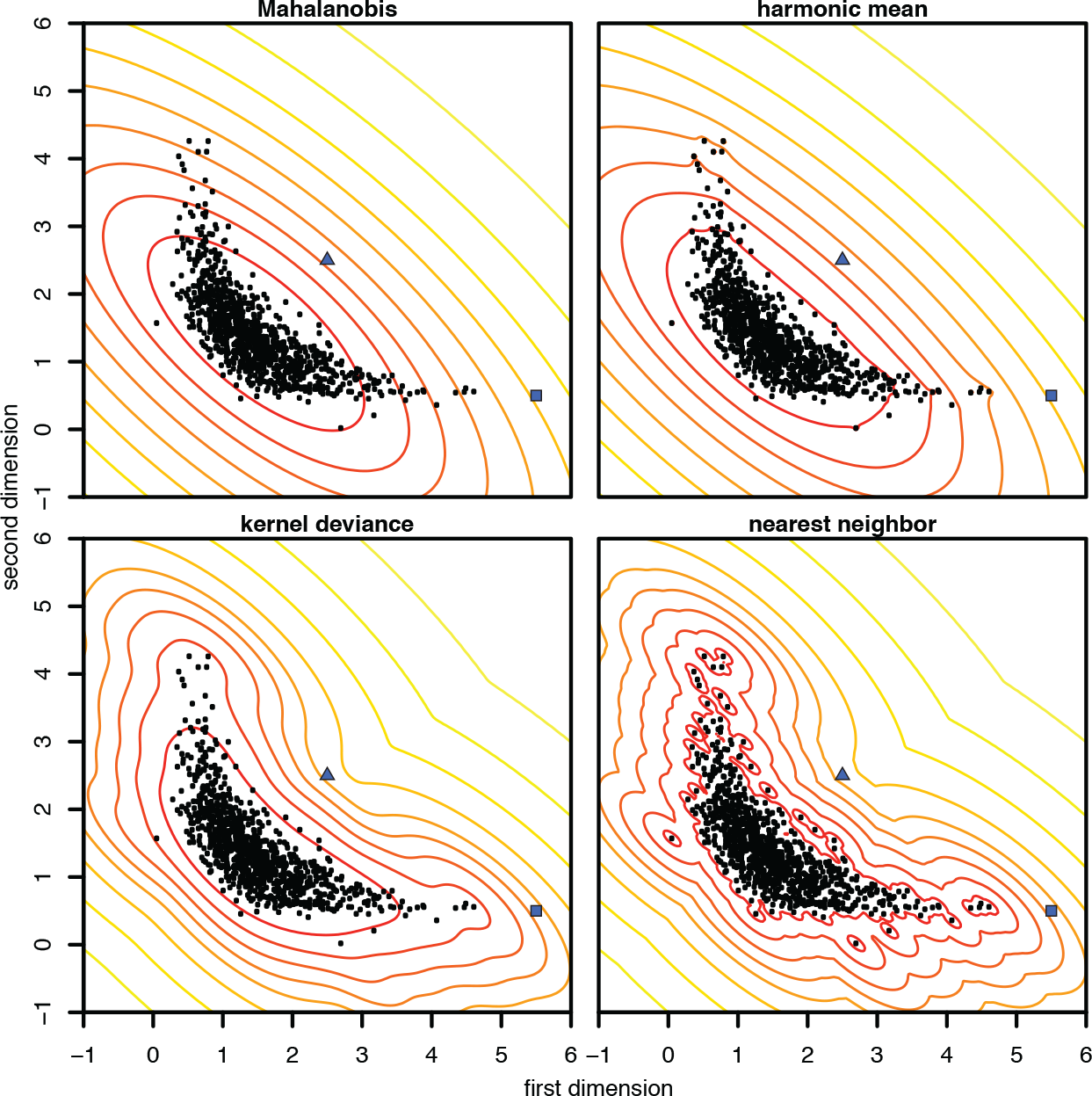
Comparison of multivariate distance measures for nonparametric example data. Black dots show the simulated data, in which the two statistics (dimensions) are assumed to follow an inverse relationship with some additional noise. Solid lines show the distance measure computed at each point in the plane, arranged in 10% quantiles (e.g. the inner ring shows the 10% of locations with the smallest distance). The blue square and triangle show particular outlier points referred to in the main text.

**Figure 4.**
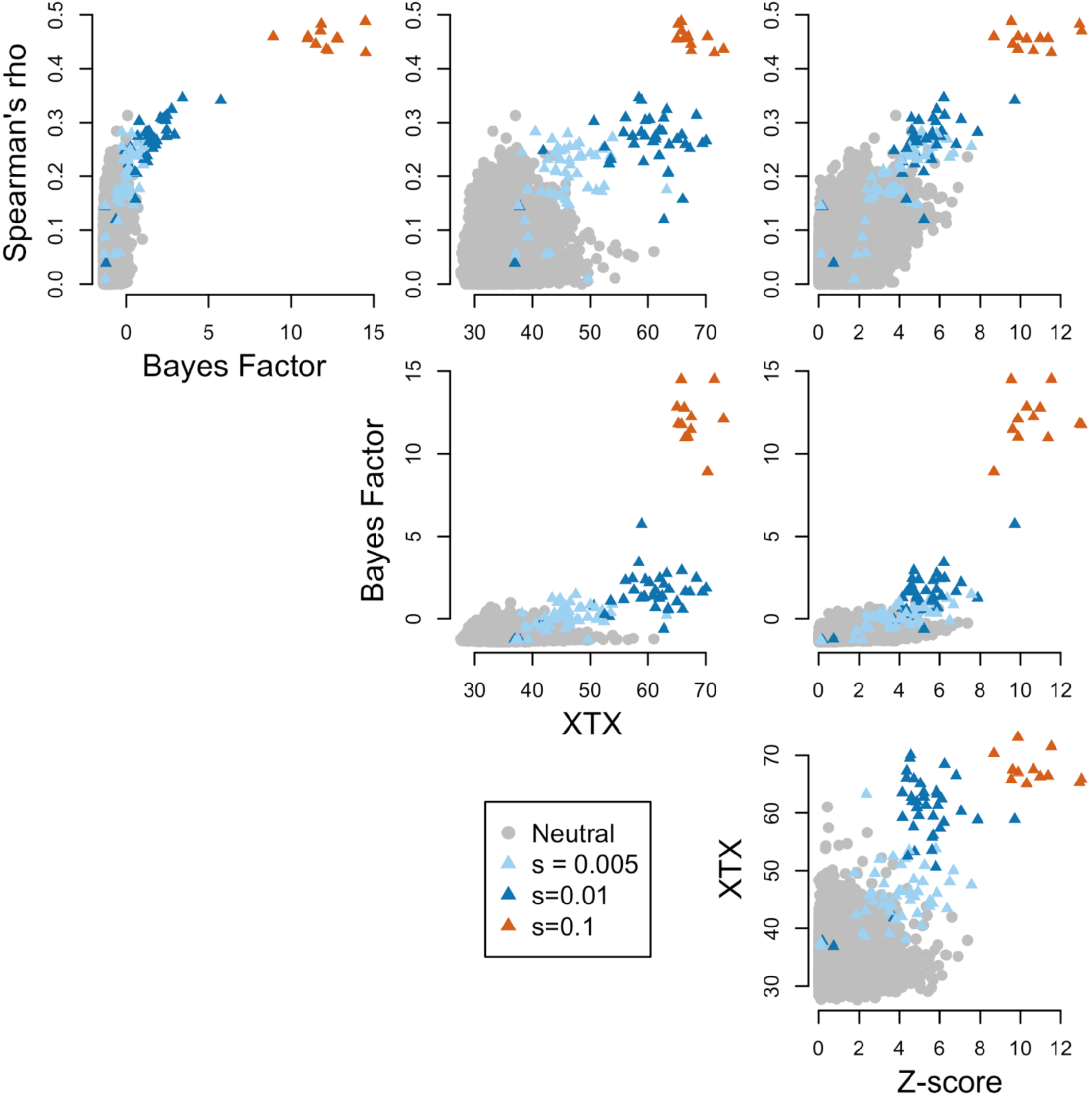
Distributions of four univariate statistics from the two refuge dataset from Lotterhos and Whitlock (2015).

*Example on simulated genomic data.* To test the power of multivariate statistics for genome scans, we applied them to a published simulated dataset that was used to test different genome scan methods (Lotterhos and Whitlock 2014, Whitlock 2015). Briefly, a landscape simulator was used to simulate haploid neutral and selected loci that adapted to an environmental cline (Lotterhos and Whitlock 2015). The landscape consisted of 360 × 360 demes and the allele frequency of each deme changed each generation according to recurrence equations for mutation, migration, selection (if applicable), and drift (Lotterhos and Whitlock 2015). For the dataset used in this example, a total of 9900 neutral and 100 selected loci (simulated under varying strengths of selection: 12 loci with s = 0.1, 38 loci with s = 0.01, and 50 loci with s = 0.005) were simulated under a two-refuge demographic expansion. Individuals were then sampled from the landscape according to the allele frequency in each deme at 30 randomly chosen locations on the landscape at 20 individuals per location. For additional details see Lotterhos and Whitlock (2014, 2015).

The simulated data were used to create a single nucleotide polymorphism (SNP) table and this data was used to perform genome scans in the programs Bayenv2 (Gunther and Coop 2013) and LFMM (Frichot et al. 2013, now implemented in the R package LEA: Frichot and Francois 2015). A total of four univariate statistics from these two programs were used in the search for multivariate outliers: (i) log-Bayes Factor (log-BF, a measure of the association between allele frequency and the environment in Bayenv2), (ii) Spearman’s rho (a measure of the association between allele frequency and the environment in Bayenv2), (iii) *X^T^X* (a measure of genetic differentiation among populations in Bayenv2), and (iv) *Z*-score (a measure of the association between genotype and the environment in LFMM). These four univariate statistics, plotted in Figure 4, were previously shown to have different strengths and weaknesses depending on sampling design and demographic history (Lotterhos and Whitlock 2015).

To illustrate the flexibility of the outlier functions implemented in MINOTAUR, we calculated multivariate outliers in two ways, corresponding to two different ways of calculating the covariance matrix *S* in equations (1) to (4). First, we used the traditional method of calculating the covariance matrix based on all the data. For high-dimensional data, estimation of the multivariate mean and covariance (location and scatter) are expected to be robust to outliers as long as the proportion of outliers in the data is less than 1/(*k*+1), where *k* is the number variables in the dataframe (Ro et al. 2015). However, we found that even in this small dataframe of only 4 variables and 10,000 loci, the 1% of selected loci (a fraction of which were true outliers) affected the estimation of the covariance matrix. For this reason, our MINOTAUR functions are designed to allow the user to input their own covariance matrix. To illustrate this use of the function, we also calculated a robust multivariate location and scatter estimate with a high breakdown point, using the ‘Fast MCD’ (Minimum Covariance Determinant) estimator with the function `CovNAMcd` in the R package `rrcovNA` (Rousseeuw et al. 1999; Todorov et al. 2011).

To compare the ability of the univariate statistics and the multivariate statistics to separate neutral from selected loci, we calculated the empirical power. The empirical power is based on using all known neutral loci to generate a null distribution, and then for each locus an empirical *p*-value is calculated based on its cumulative frequency in this null distribution. To control for false discovery rate, empirical p-values were converted to q-values (in the R package qvalue: Dabney and Storey 2014) and loci with a *q*-value less than 0.05 were retained as positive hits (a *q*-value of 0.05 has a desired rate of 5 false positives out of 100 positive hits).

For the univariate statistics, the empirical power was highest for log-BF (0.54) and lowest for *Z*-score (0.15), with Spearman’s rho (0.46) and *X^T^X* (0.42) also showing moderate power. For the multivariate statistics with the default covariance estimation, the empirical power was high for harmonic mean distance and Mahalanobis distance (0.41 for both), with kernel density and nearest neighbor distance performing poorly in this case (0.09 for both) (Supplementary Figure S2). For the user-input covariance matrix estimated with a high breakdown point (i.e., less influenced by outliers), the empirical power was highest for harmonic mean distance and Mahalanobis distance (0.58 for both), with kernel density and nearest neighbor distance still performing poorly (Figure 5). This final example illustrates the potential of Mahalanobis and harmonic mean distance to improve the signal-to-noise ratio in genome scans, because the empirical power in this case was higher than any univariate statistic alone.

**Figure 5.**
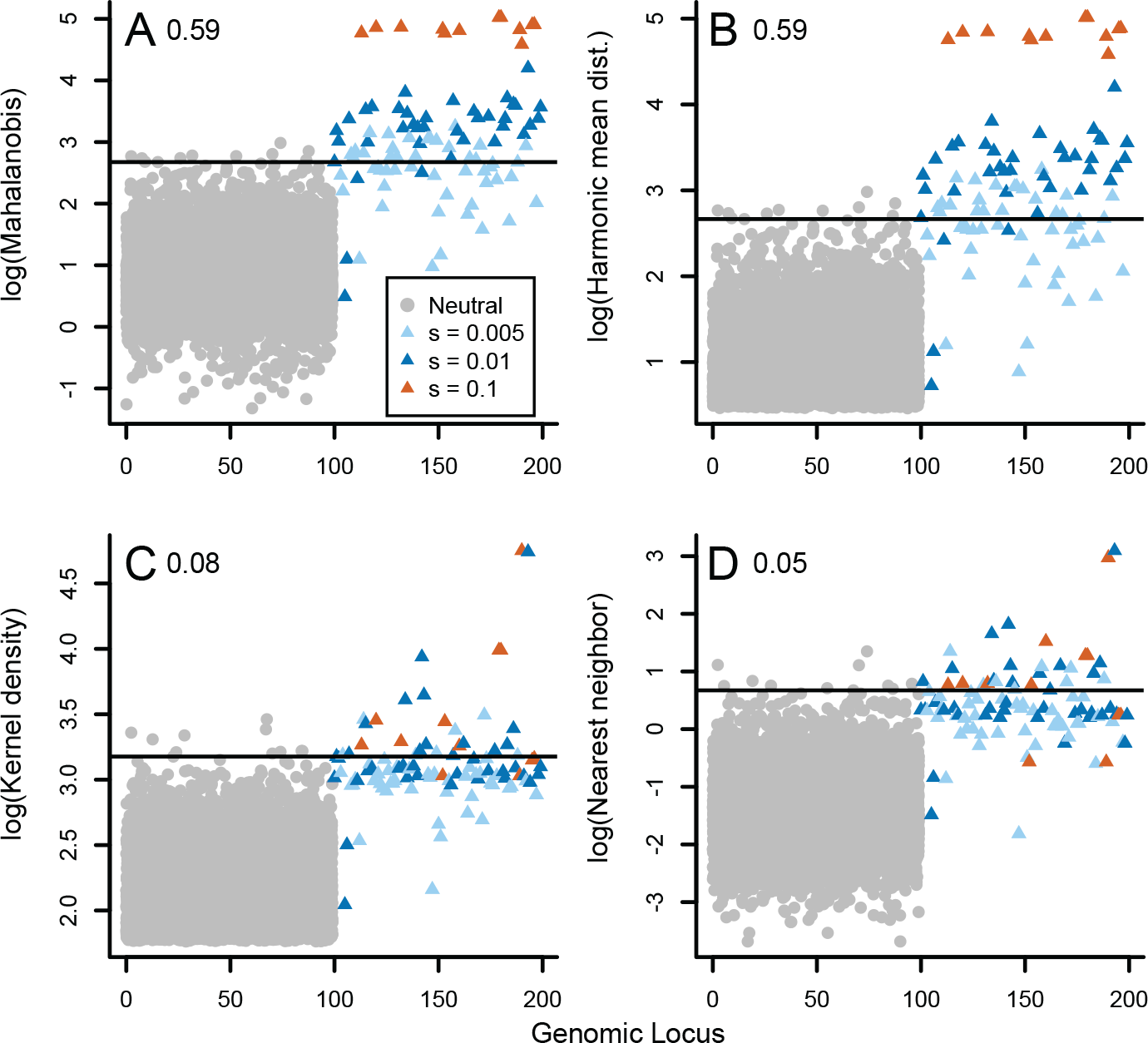
Distributions of the four multivariate compound statistics applied to the four univariate statistics shown in Figure 2. The MCD calculation of the covariance matrix was used. All 9900 neutral loci are plotted on indexes 0-100, and the selected loci are plotted on indexes 100-200. Note log transformation of each variable on the y-axis for: A) Mahalanobis distance, B) Harmonic mean distance, C) Kernel density, and D) Nearest Neighbor distance. The empirical power of the statistic to discriminate neutral from selected loci (see main text for details) is shown in the upper left hand corner.

## Discussion

Although the number of packages for population genetic data analysis in the R software is rapidly increasing (http://popgen.nescent.org/PACKAGES.html), basic tools for manipulating and visualizing genome-scale datasets have so far been lacking. MINOTAUR fills this gap using the R Shiny Dashboard package to implement a GUI that makes it easy to upload, manipulate, analyze, and visualize genomic data.

The multivariate metrics calculated in MINOTAUR contribute to a growing number of multivariate tools implemented in the R environment (see Supplementary Table S1). Methods that are influenced heavily by the distance of a point from the centroid in multivariate space (such as Mahalanobis and the harmonic mean distance) will perform differently compared with methods that are influenced mainly by the sparseness of points in multivariate space (such as nearest neighbor distance and kernel density), as illustrated in the examples here. However, depending on how the data are distributed, the harmonic mean distance may be influenced by both these factors. For a single simulated dataset, we found that robust use of the Mahalanobis or harmonic mean distance (i.e., when the covariance matrix used was estimated with a high breakdown point) could have higher power than any single univariate statistic alone. Although nearest neighbor distance and kernel density deviance performed poorly on the simulated genomic data, they may be useful in application to other kinds of nonparametric data, as illustrated in our examples (Figures 3 and S1). Further evaluation, however, will be needed on both simulated and empirical data to determine whether multivariate outlier approaches will improve the signal-to-noise ratio in genome scans.

The MINOTAUR package is designed to complement existing tools for the analysis and integration of genome-scan data. Thus, in addition to providing its own tools for genome-scale analyses, MINOTAUR can serve as a platform for the further analysis and visualization of results generated by other R packages. Examples include results from differential gene expression (LIMMA: Ritchie et al. 2015; DESeq: Anders and Huber 2010; SeqGSEA: Wang and Cairns 2014), outliers for genetic differentiation (OutFLANK: Whitlock and Lotterhos 2015; PCAdapt: Luu and Blum 2015), genetic-environment associations (LEA: Frichot and François 2015), or genome-wide association studies (e.g. GenABEL: Aulchenko et al. 2007; BlueSNP: Huang et al. 2013).

Recent developments such as the R Shiny and Shiny Dashboard environments (Chang 2015; Chang et al. 2016) dramatically aid in the development of R-based user-friendly web interfaces. Taking advantage of these tools, MINOTAUR is able to offer a new platform for visualizing and integrating genomic data that may appeal to molecular ecologists, modellers, statisticians, and public health agencies.

## Resources

**Availability:** Upon acceptance for publication, MINOTAUR will be distributed on CRAN (http://cran.r-project.org/) and be available for R on Windows, Mac OSX, and Linux platforms. Currently, MINOTAUR can be accessed via the following steps:

- `install.packages(“devtools”, dependencies=TRUE)`
- `library(devtools)`
- `install_github(“NESCent/MINOTAUR”, build_vignettes=TRUE)`
- `library(MINOTAUR)`
- `MINOTAUR()`

Note to reviewers: If you are facing issues with installation, try updating to the newest version of R and reinstalling devtools from source. MINOTAUR has been tested on R version 3.3.0.

**Licence:** GNU General Public Licence (GPL) >= 2.

**Documentation**: Besides the usual package documentation, MINOTAUR is released with a tutorial which can be opened by typing: vignette(“MINOTAUR“).

**Development:** The development of MINOTAUR is hosted on GitHub: (https://github.com/NESCent/MINOTAUR).

## Acknowledgments

The resource reported in this paper began at the Population Genetics in R Hackathon, which was held in March 2015 at the National Evolutionary Synthesis Center (NESCent) in Durham, NC, with the goal of addressing interoperability, scalability, and workflow building challenges for the population genetics package ecosystem in R. The authors were participants in the hackathon, and are indebted to NESCent (NSF #EF-0905606) for hosting and supporting the event. RV, CC, DCC, SMS benefited from travel support to attend this Hackathon.

### Author Contributions

RV and KEL conceptualized the study. RV derived the compound outlier measures and implemented them in R. RV and CC conceptualized and implemented the Shiny Dashboard GUI. CC managed R package development, package structure and documentation. All authors contributed R code for plotting in the Shiny Dashboard. DCC wrote the vignette for the package and estimated function computational time. KEL performed analysis of the performance of compound measures on simulated data. All authors contributed to writing this manuscript.

---

**Table S1.** Table of multivariate outlier statistics in other R packages that could be used in the context of genomic scans.

| Multivariate Outlier Statistic | R Package | R Function | Brief Description | Reference |
| --- | --- | --- | --- | --- |
| Hierarchical Clustering Ranks | DMwR | outliers.ranking() | Uses an agglomerative hierarchical clustering algorithm to rank outlierness. | Torgo 2011 |
| Projection Congruent Subset | FastPCS | FastPCS() | Computes fast and robust multivariate outlyingness index. | Vakili & Schmitt 2014 |
| Kernel Density Estimator | ks | kde() | Computes the kernel density estimate for up to 6 dimensional datasets. | Duong 2007 |
| Mahalanobis Distance | mvoutlier | locoutNeighbor() | Computes global and pairwise Mahalanobis distances for outlier visualization with number of neighbors varying and fraction of neighbors fixed. | Filzmoser & Gschwandtner 2015 |
| Mahalanobis Distance | mvoutlier | locoutSort() | Computes global and pairwise Mahalanobis distances for interactive outlier visualization. | “ |
| Mahalanobis Distance | mvoutlier | locoutPercent() | Computes global and pairwise Mahalanobis distances for outlier visualization with number of neighbors fixed and varying fraction of neighbors. | “ |
| Principal Components Distance | mvoutlier | pcout() | Principal components distances are used to identify weighted location and scatter of outliers. | “ |
| Mahalanobis Distance | mvoutlier | sign1() | Principal components are used to calculate Mahalanobis distance covariance matrix and a critical value cutoff is used to determine outliers from chi-squared distribution. | “ |
| Principal Components Distance | mvoutlier | sign2() | Principal components distances are computed and transformed to approach a chi-squared distribution and a critical value cutoff is used to detect outlier. | “ |
| Adjusted Mahalanobis Distance | mvoutlier | arw() | Adjusts outlier rejection thresholds by using an adaptive reweighting estimator and determines outliers by the supremum of the difference between Mahalanobis distance and the theoretical distribution function. | “ |

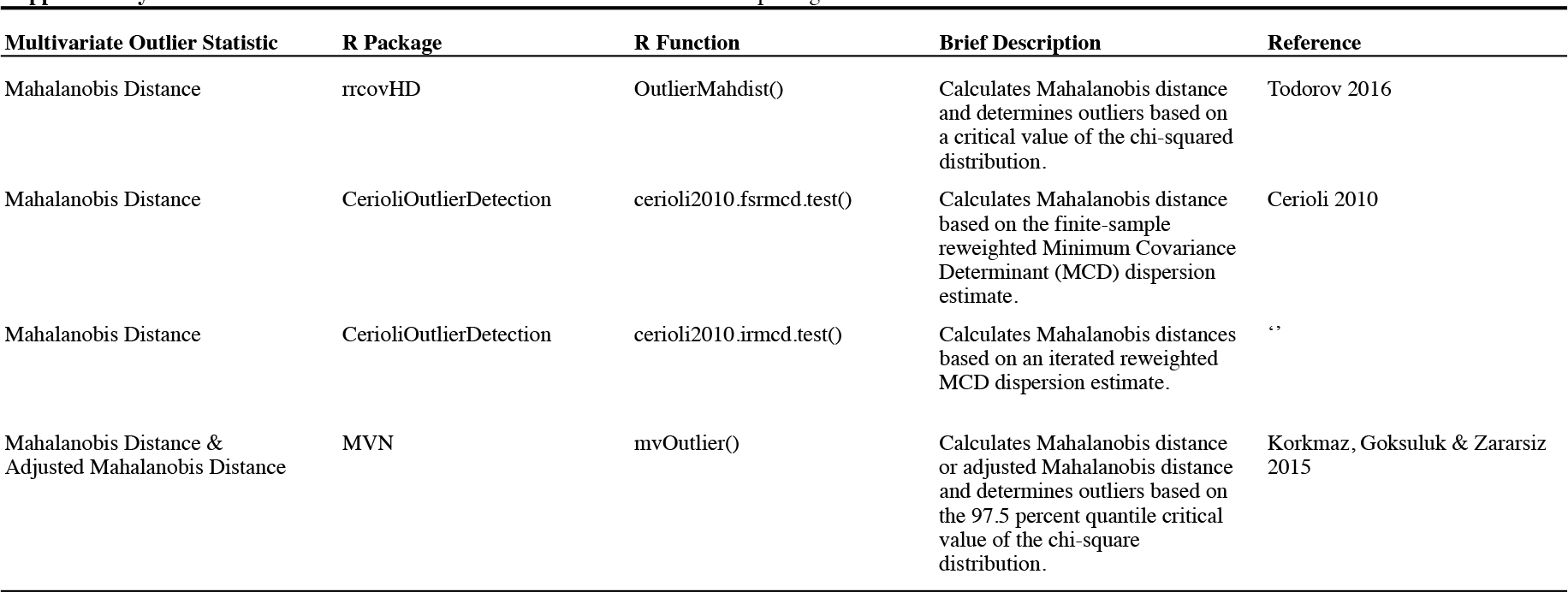

**Table S2.** Computation times for the four multivariate outlier detection methods in MINOTAUR for datasets up to 100,000 loci (rows) and 20 variables (columns) in hh:mm:ss.ms format. Run times were determined using an Apple iMac with a 3.1 GHz Intel Core i5 processor and 32 GB of RAM running Apple OSX 10.9.5 and R version 3.2.3. Note that the kernel density deviance includes both the maximum likelihood estimation of the optimal bandwidth and the density calculations based on the optimal bandwidth.

| No. Loci | No. Variables | Mahalanobis distance | Harmonic mean distance | Kernel density deviance | Nearest neighbor distance |
| --- | --- | --- | --- | --- | --- |
| 1000 | 5 | 00:00:00.001 | 00:00:00.040 | 00:00:01.233 | 00:00:00.034 |
| 1000 | 10 | 00:00:00.002 | 00:00:00.098 | 00:00:02.366 | 00:00:00.094 |
| 1000 | 15 | 00:00:00.003 | 00:00:00.188 | 00:00:04.303 | 00:00:00.185 |
| 1000 | 20 | 00:00:00.005 | 00:00:00.318 | 00:00:07.548 | 00:00:00.317 |
| 5000 | 5 | 00:00:00.003 | 00:00:00.986 | 00:00:30.215 | 00:00:00.829 |
| 5000 | 10 | 00:00:00.008 | 00:00:02.431 | 00:00:57.794 | 00:00:02.382 |
| 5000 | 15 | 00:00:00.014 | 00:00:04.809 | 00:01:49.900 | 00:00:04.622 |
| 5000 | 20 | 00:00:00.024 | 00:00:07.968 | 00:02:46.393 | 00:00:07.821 |
| 10000 | 5 | 00:00:00.006 | 00:00:03.922 | 00:02:00.694 | 00:00:03.328 |
| 10000 | 10 | 00:00:00.017 | 00:00:09.728 | 00:03:52.081 | 00:00:09.310 |
| 10000 | 15 | 00:00:00.169 | 00:00:18.773 | 00:06:53.011 | 00:00:18.487 |
| 10000 | 20 | 00:00:00.041 | 00:00:32.415 | 00:11:10.482 | 00:00:31.735 |
| 50000 | 5 | 00:00:00.045 | 00:01:37.808 | 00:50:19.078 | 00:01:24.120 |
| 50000 | 10 | 00:00:00.095 | 00:04:13.621 | 01:40:03.603 | 00:04:07.027 |
| 50000 | 15 | 00:00:00.161 | 00:09:12.221 | 00:19:36.847 | 00:08:51.240 |
| 50000 | 20 | 00:00:00.240 | 00:16:38.589 | 05:45:21.387 | 00:16:12.965 |
| 100000 | 5 | 00:00:00.085 | 00:06:35.265 | 03:21:34.758 | 00:05:35.996 |
| 100000 | 10 | 00:00:00.184 | 00:17:01.708 | 09:28:30.378 | 00:16:24.535 |
| 100000 | 15 | 00:00:00.324 | 00:36:19.473 | 13:28:52.158 | 00:37:20.698 |
| 100000 | 20 | 00:00:00.487 | 01:14:47.783 | 24:45:25.372 | 01:14:24.686 |

**Figure S1.**
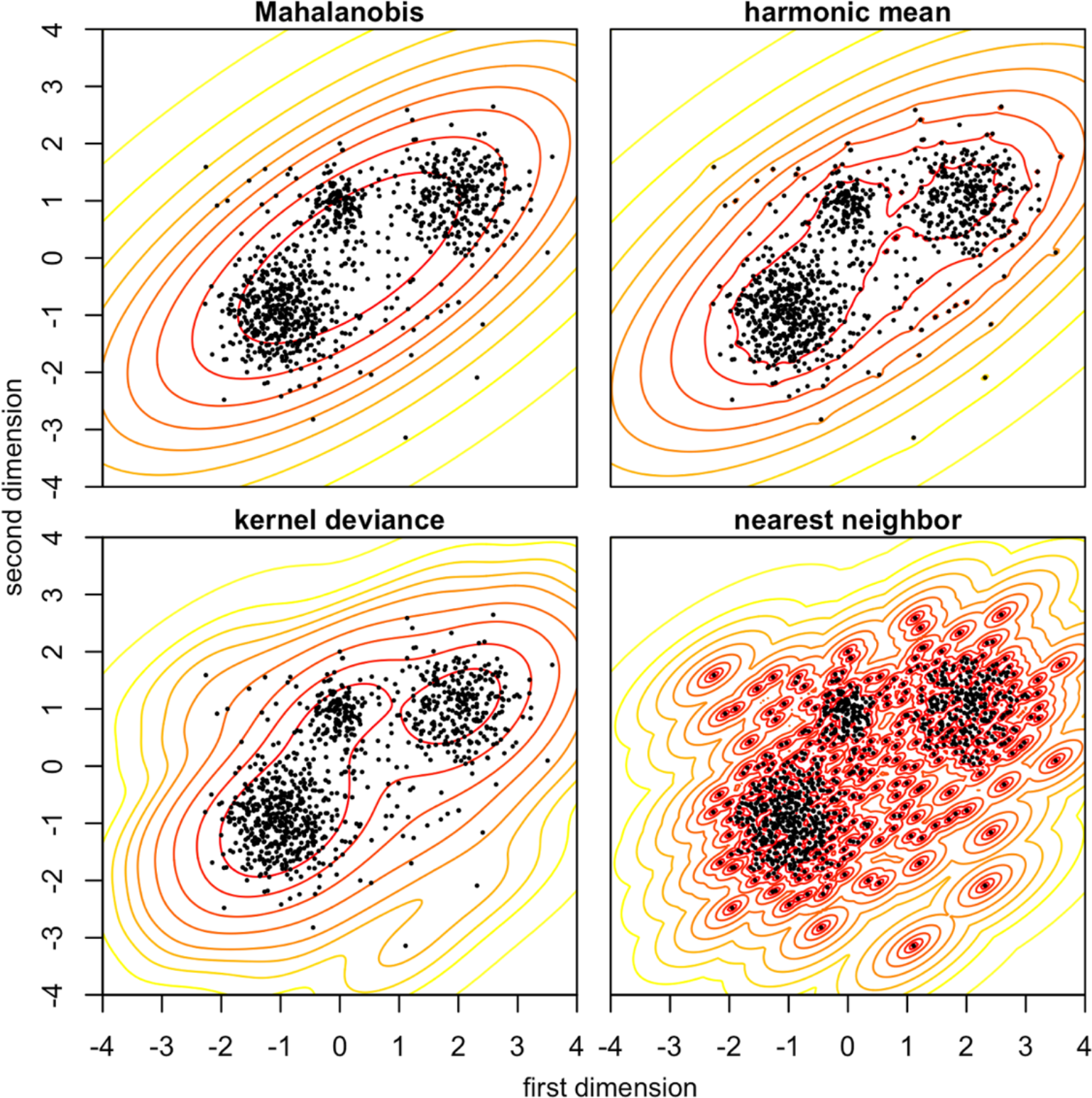
Comparison of multivariate distance measures for multi-modal example data. Black dots show the simulated data, drawn from a bivariate normal mixture model. Solid lines show the distance measure computed at each point in the plane, arranged in 10% quantiles, equivalently to Figure 3 in the main paper.

**Figure S2.**
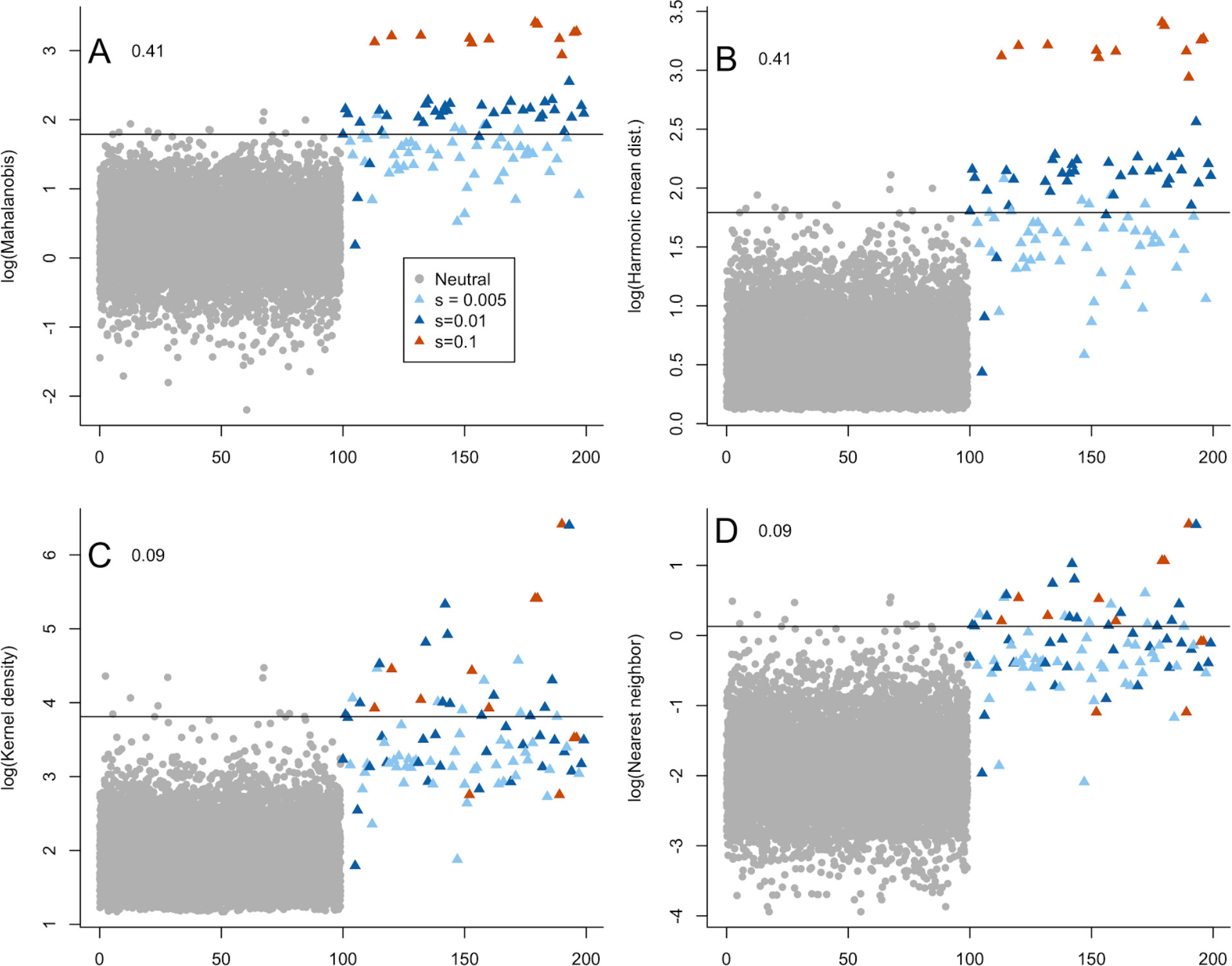
Analogue to Figure 5 in the main paper, but with a default estimate of covariance using all the data.

